# Fatigue, Alertness and Risk Prediction for Shift Workers

**DOI:** 10.1101/2021.01.13.426509

**Authors:** Sean F. Cleator, Louise V. Coutts, Robert Phillips, Ruth Turner, Derk-Jan Dijk, Anne Skeldon

## Abstract

**Executive summary:**

1. This report describes the principal outcomes of an Impact Acceleration Account project (grant number EP/I000992/1) between the University of Surrey and Transport for London carried out between Oct. 2019 and Mar. 2020.
2. The aim of the project was to compare the Health and Safety Executive (HSE) Fatigue Risk tool with SAFTE and other more recent models of fatigue, where fatigue here primarily means a reduced ability to function effectively and efficiently as a result of inadequate sleep.
3. We have not sought to discuss the useability of the HSE Fatigue Risk tool or SAFTE since this has been discussed comprehensively elsewhere (e.g. [1, 2]). We have instead focussed on the fundamental principles underlying the models.
4. All current biomathematical models have limitations and make asumptions that are not always evident from the accompanying documentation. Since full details of the HSE Fatigue Risk tool and the SAFTE model are not publicly available, Sections 1 and 2 give a mathematical description of the equations that we believe underlie each of these models.
5. A comparison of predictions made by our versions of the HSE and SAFTE equations for one particular shift schedule of relevance to the UK and global tunnelling and construction industries is shown in Section 3. In this comparison, we use data collected durings TfL’s Crossrail project by Dragados^1^. Essentially, both models give broadly the same message for the schedule we looked at, but the ability to display fatigue as it develops within a shift is a strength of SAFTE.
6. A summary of the strengths and limitations of the use of these kind of scheduling tools is given in the final Section 4. Limitations include:
  - Models do not describe fatigue during times when people are not in shift (e.g. driving home). However, they could readily be extended to do so.
  - Models assume people start well-rested. This is not always a good assumption and can lead to an under-estimate of fatigue.
  - Most models are currently based on population averages, but there are large individual different. It would be possible to further develop models to include uncertainty in fatigue predictions associated with individual differences.
  - Few mdels include the light environment, which is important both to promote short-term alertness and facilitate circadian alignment.
  - Models are not transparent, which makes them hard to independently validate.
  - It is hard to relate the outputs of current models to measureable outcomes in the field.
7. We also discuss briefly recent developments in mathematical modelling of fatigue and possible future directions. These include
  - Guidance on scheduling and education on sleep and fatigue should be considered at least as important as current biomathematical models.
  - Only by analysing and integrating high quality individual data on sleep, fatigue, performance, near misses, accidents, actual shift patterns with models can we develop better models and management systems to reduce fatigue and associated risks. Wearables combined with apps present a great opportunity to collect data at scale but need to be used appropriately.
  - The importance of making time for sleep is not always recognised. Education, early diagnosis of sleep disorders such as sleep apnea, and self-monitoring all have a role to play in reducing fatigue-related risk in the work-place.
8. Section 3 and Section 4 may be understood without reading the intermediate more mathematical sections.

## 1 The HSE Fatigue and Risk Tool: Fatigue Index

### 1.1 Background

The HSE Fatigue and Risk tool [3] calculates two quantities, the Fatigue Index and the Risk Index, aimed at providing guidance on the fatigue associated with any specified shift schedule. Here we focus on the Fatigue Index. We will be making extensive reference to [3], so from this point will refer to it as the QuinetiQ research report.

The QuinetiQ research report states that the Fatigue Index represents the probability that someone will score highly (eight or nine) on the Karolinska Sleepiness Scale (KSS), a nine-point scale ranging from one (extremely alert) to nine (extremely sleepy and fighting sleep).

The Fatigue Index considers sleepiness to arise from the interaction of three components:

(i) a cumulative component, 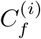, which models the effect of accumulated sleep debt; (ii) a shift^2^ timing component, 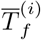, which takes into account the timing of the shift; (iii) a job type/breaks component, 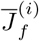, which allows for different types of job to lead to different levels of sleepiness and takes into account breaks.

The Fatigue Index for a shift on day *i*^3^, *FI*^(*i*)^, is given by

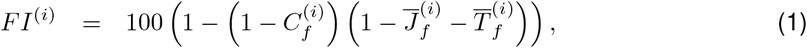

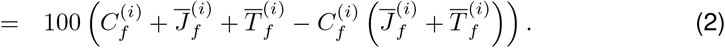

The three components are scaled and each takes a value up to a maximum of approximately one third. ^4^.

The basis for this formula is not clear from the documentation, but it can be seen from equation (2) that the three components are essentially additive with a nonlinear correction term that downweights the job type and duty timing components when cumulative fatigue is high. Multiple factors feed into the cumulative component, the shift timing component and job type component. A schematic giving the overall steps is shown in Fig. 1. The formula are discussed below.

**Figure 1:**
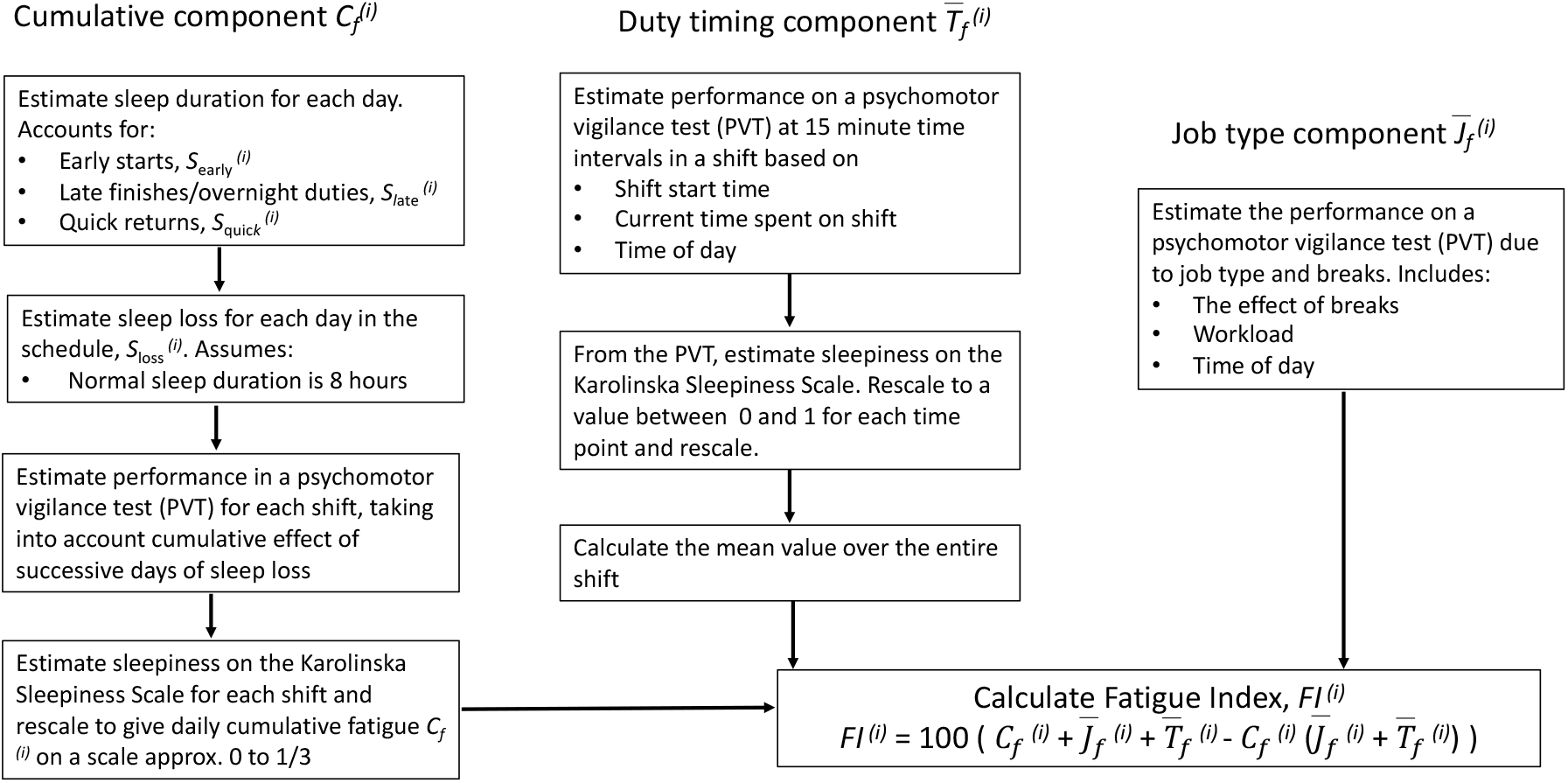
Schematic summarising the process for calculating the Fatigue Index.

### 1.2 Cumulative component, 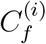

The cumulative component is based on sleep debt and, as indicated in Fig. 1, is calculated by first estimating the sleep duration for each day. This is then used to calculate the sleep loss. Sleep loss for each day is then used to estimate a baseline performance value on a psychomotor vigilance task (PVT).

This baseline PVT value is then adjusted up or down depending on performance the previous day. This scaling up or down based on prior performance in effect models the impact of accumulated sleep loss. For shifts starting during the day, defined as shifts starting between 03:00 and 15:00, there is an additional small adjustment based on start time.

There is a final rescaling step that rescales PVT to values that relate to the Karolinska Sleepiness Scale (KSS, a scale from one to nine) and then to a value between zero and approximately one third.

Each of these steps is discussed in more detail below.

#### 1.2.1 Daily sleep loss, 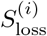

The HSE tool assumes that a normal night’s sleep is eight hours and that shorter than normal sleep duration may occur as a result of three factors: an early shift start; a late shift finish or overnight shift, or as a result of two successive shifts are close together (a quick return). Specifically, sleep duration 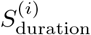 on day *i* is

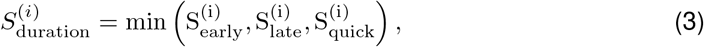

where 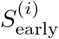 is the sleep duration due to an early start in the morning; 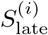 is the sleep duration due to a late finish in the evening or an overnight shift, and 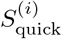 is the sleep duration as a result of a quick return. All sleep durations are measured in hours and refer to the sleep duration on the day leading up to the shift on day *i*. ^5^

Consequently, sleep loss, 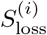, in hours for the night leading up to day *i* is modelled as

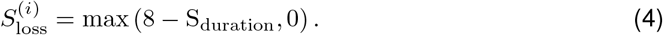

In the QuinetiQ report, the formula for 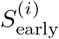 is described as a quadratic fit to data from train drivers showing mean sleep duration as a result of an early start. These data show that start times of 04:30 results in only 5 h of sleep prior to the shift. With each 1 h delay in start time there is approximately 0.5 h of additional sleep, until by a start time of 09:30, almost 7.5 h of sleep are obtained. We have refitted these data, giving,

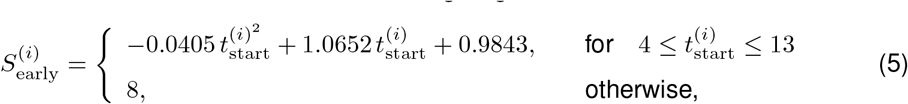

where 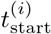 is the start time of the shift on day *i* in hours on a 24 h clock. The original data is shown in Fig. 2(a) on which we have superimposed equations (5).

**Figure 2:**
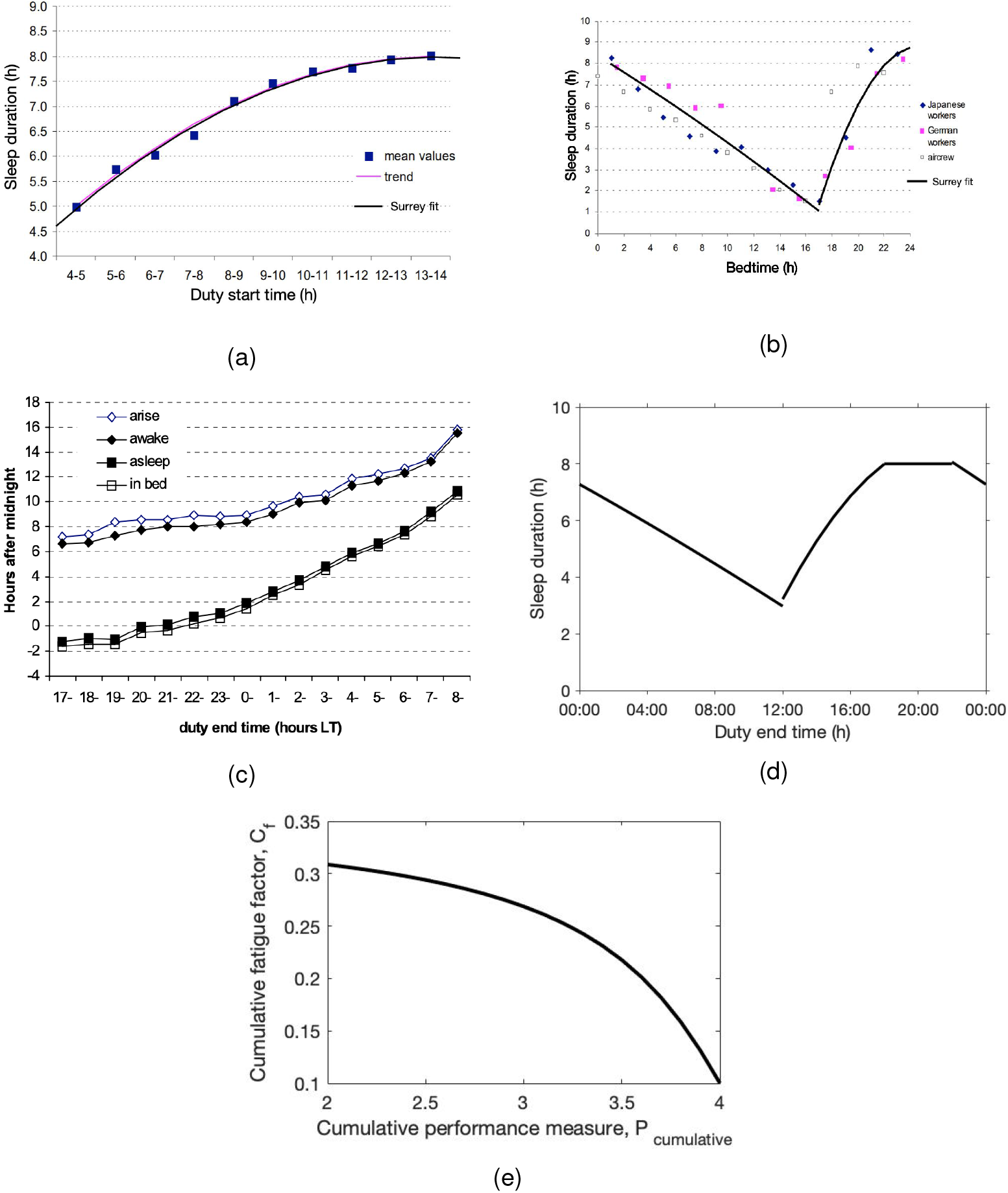
(a) Expected sleep duration for early shift starts. Equations (5) superimposed on Fig. C2 from the QuinetiQ report [3]. (b) Expected sleep duration as a result of bedtime. Equations (6) superimposed on Fig. C3 from the QuinetiQ report [3]. (c) Bedtime related to shift end time, Fig. 24 from [4]. (d) Sleep duration as a function of shift end time, as given by equations (7). (e) Cumulative fatigue component *C*_*f*_ as a function of cumulative performance *P*_cumulative_, as given by the sequence of transformations in equations (11)-(14). Panels (a) and (b) have been included with permission from the Health and Safety Executive (HSE). The original figures and report can be found here https://www.hse.gov.uk/research/rrhtm/rr446.htm. Panel (c) is reproduced with permission of the Civil Aviation Authority (CAA).

In the QuinetiQ report, 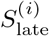, is described as being derived from the data on bedtimes for Japanese and German workers, reproduced in Fig. 2(b). These data show that going to bed at midnight results in 8 h sleep. However, sleep duration decreases approximately linearly with bedtime, with a bedtime of 10:00 resulting in approxmiately 4 h sleep and a bedtime of 14:00 resulting in only 2 h sleep. The pattern changes after 17:00, with sleep duration increasing as bedtimes approach ‘normal’ bedtimes. Refitting to these data gives sleep duration based on bedtime, 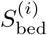, on day *i* as

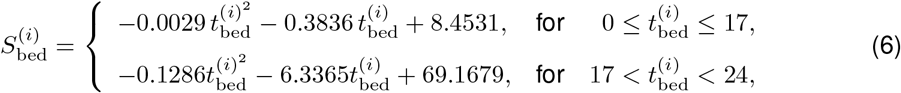

where 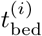 is the bedtime on day *i* in hours on a 24 h clock. Equations (6) have been superimposed on the original data in Fig. 2(b).

In order to deduce bedtime from shift end time, it is stated in the QuinetiQ report that data from an internal QuinetiQ report was used resulting in a scale that gives a minimum sleep duration of 3 h for shifts ending at 12:00, and 8 h (no sleep loss) for shifts ending at 18:00. We could not access the QuinetiQ report, although some indication of how shift time and bedtime are related can be deduced from Fig. 20 from [4], reproduced in Fig. 2(c). Primarily based on the description that there is a minimum of sleep duration of 3 h for shifts ending at 12:00 and no sleep loss for shifts ending at 18:00, we have assumed a piecewise scaling of the fit given in equation (6), giving

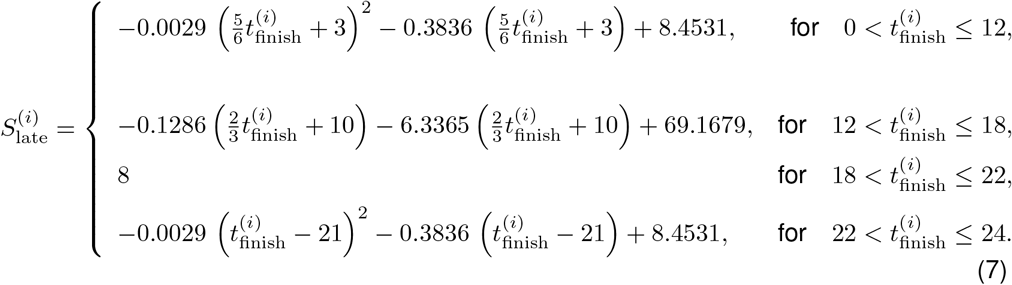

The resultant dependence of sleep duration on shift end time is shown in Fig. 2(d).

Note the piecewise scaling effectively says that if a shift finishes at midnight, it takes 3 h between the end of the shift and bedtime, but this gap between end of shift and bedtime decreases linearly to 1 h for shifts that end at 12:00. For shifts that end after 12:00, the time between the end of the shift and bedtime increases until for shifts ending at 18:00, bedtime is 4 h later, at 22:00. This may seem early, but is consistent with the data shown in Fig. 2(c) and it should be noted is intended to model late finishes or overnight duties.

In the Quinetiq report, sleep duration, 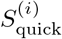, as a consequence of quick returns is given as

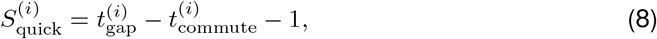

where 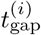 is the time between the shift on day *i* and the shift on day 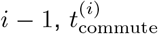 is the commute time and it is assumed that one further hour is needed for personal needs.

#### 1.2.2 Using sleep loss to calculate performance 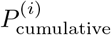

Sleep loss has been related to longer reaction times in a performance vigilance task (PVT). Based on laboratory data, the HSE fatigue tool calculates a baseline performance, 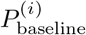 on each day based on the sleep loss in the previous 24 h. Specifically,

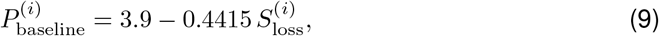

i.e. performance decreases linearly with sleep loss. This baseline value for each day is then updated depending on whether or not the predicted baseline performance is better or worse than the day before. Specifically,

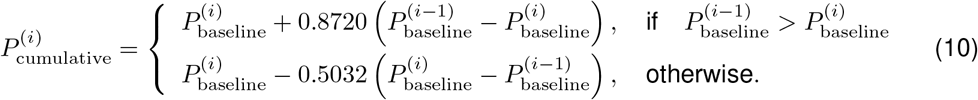

There are two further adjustments made (i) a small circadian adjustment made to shifts with start times between 03:00 and 15:00 (ii) an adjustment if the gap between two shifts is small to take into account split shifts.

#### 1.2.3 Calculating the cumulative component, 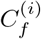 based on performance

Finally, the daily cumulative component of the Fatigue Index, 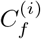 is calculated from the cumulative estimate of performance. Specifically, first scaling the PVT measure to the seven-point Samn Perelli sleepiness scale,

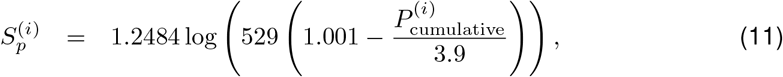

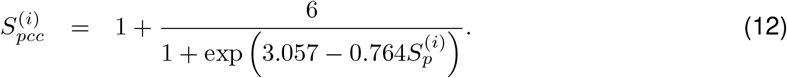

Subsequently, a further rescaling to a measure related to the Karolinska Sleepiness Scale (KSS).

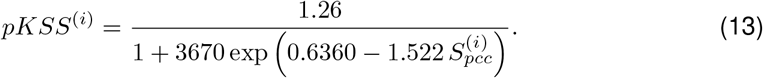

It is not clear where these various scalings come from. This is a critical step to understanding the description of the output of the tool as a measure of the probability of scoring eight or nine on the KSS.

Finally, the cumulative fatigue component, 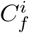 is calculated,

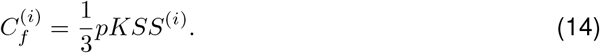

The net effect of these various transformations scales the performance in a nonlinear way to the cumulative fatigue component, as shown in Fig. 2(e). Much of the nonlinearity is a result of the nonlinear scaling from PVT to sleepiness measures, as given by equation (11). The subsequent scalings close to linear. From Fig. 2(e) it can be seen that the maximum value of 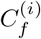 is approximately one third.

### 1.3 Shift timing component, 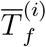

In the HSE Fatigue Index, three factors are used to calculate the effect of shift timing on fatigue: the start time of the shift; the length of the shift, and the time of day throughout the shift. The effect on fatigue of these three factors is considered to be additive, so at any time point *t*.

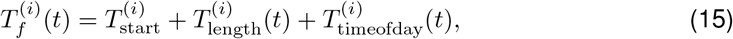

where 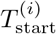 is the contribution due to start time, 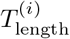 is the contribution due to the length of the shift and 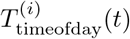 is the contribution due to time of day.

Since the shift timing component changes throughout the shift, to calculate a single value for each shift, 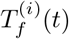 is calculated for every 15 minutes during the shift and then the mean over all the time points found to give 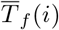.

The equation for 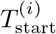 is described in the QuinetiQ report as a daily oscillation and the amplitude and acrophase (time of the maximum) are given. Specifically

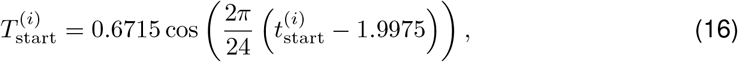

where 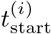 is the shift start time in hours.

The component of fatigue due to the time of day through the shift is modelled using a cosine, with a maximum at 05:15,

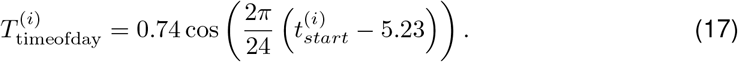

The component due to the length of the shift period increases linearly with time through the shift,

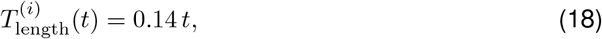

where *t* is time in hours. We note that the factor 0.14 is based on a commute time of approximately 30 minutes.

### 1.4 Fatigue: Job type / breaks component, 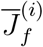

The QuinetiQ report states that the increase in fatigue as a result of continuous activity is modelled as a negative exponential function, with the more intense the activity the faster the increase. When breaks occur, there is some recovery. A short break (e.g. two minutes) temporarily halts the increase. If it is long (e.g. 30 minutes), fatigue returns to baseline, with 50% recovery after approximately 15 minutes.

The description given would give a time course of fatigue across the shift from which an average for the shift would need to be calculated. Note that the effect of job type as described should be independent of the shift start time. However, this is not consistent with outputs of the tool which suggest that the modelling of 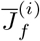 also includes a time of day effect.

### 1.5 Example: comparison of 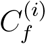 from equation (14) with outputs from the HSE tool

To demonstrate that the formula for 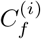 within this report give a reasonable approximation to the HSE Fatigue Index we show results for a ‘7473’ schedule. Our motivation is not to replicate the HSE tool, but to make sure that we understand the theoretical basis for the underlying computations.

The 7473 schedule is a 21 day repeating shift pattern of seven days working 07:00–19:00, four days rest, seven nights working 19:00–07:00, followed by three days rest, as shown in Fig. 3(a). Shift timing is indicated by the coloured horizontal bars. Successive days are plotted from top to bottom. The width of the plot covers two days, so that the top row shows the shift on day one followed by the shift on day two. On the next row, the shift on day two is re-plotted and followed by the shift on day three. The reason for double-plotting in this way is so that both shifts that occur during the day and shifts that occur during the night can be seen clearly.

**Figure 3:**
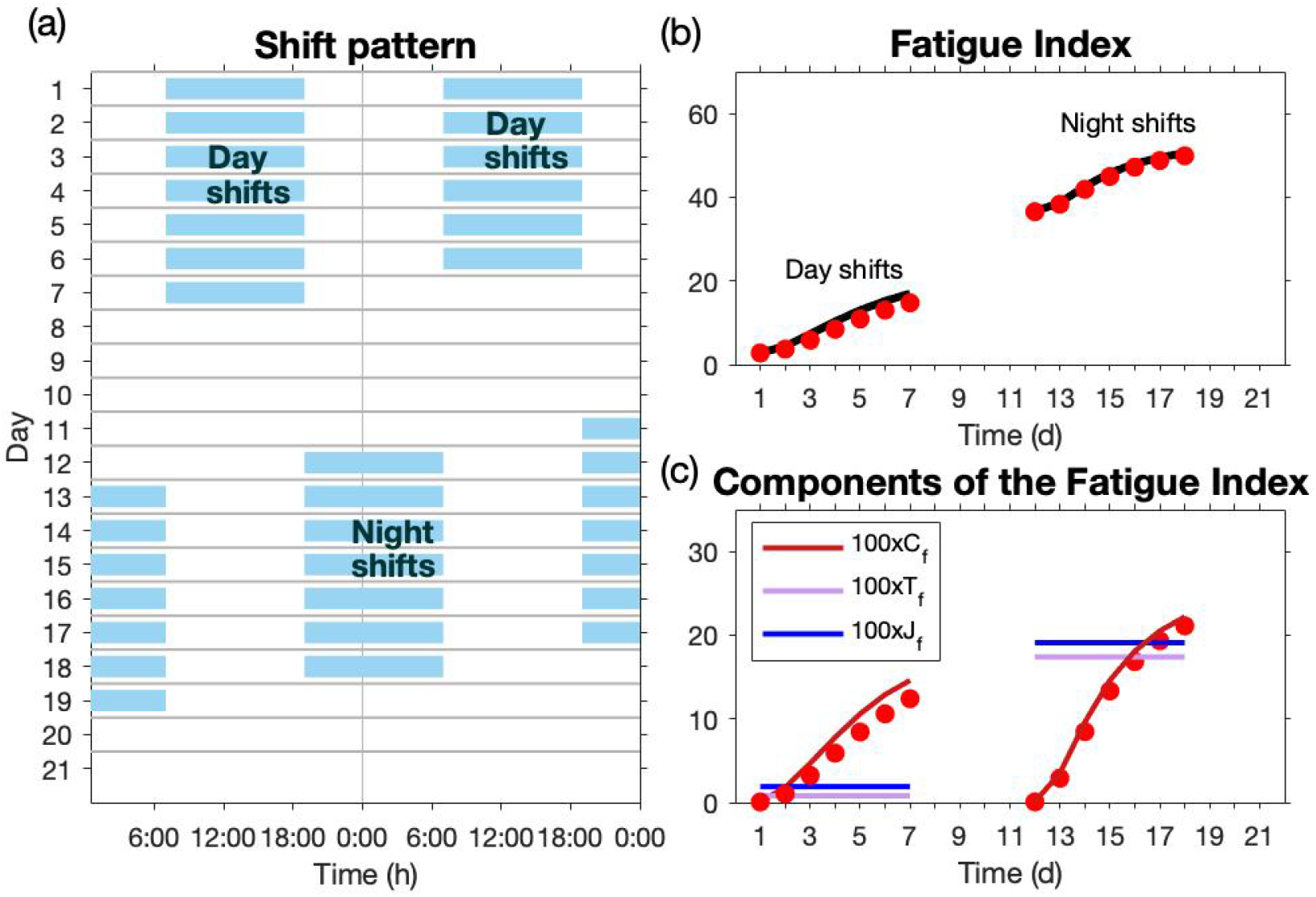
(a) Double-plot of the 7473 shift pattern. (b) Fatigue Index, *FI* from the HSE tool (black line) and from our calculation (red dots). (c) Cumulative, shift timing and job type components from the HSE tool with the cumulative component from equation (14) multiplied by 100 to be consistent with the output of the tool (red dots).

The HSE tool was used with the following defaults: commuting time: 40 minutes, workload: moderately demanding, little spare capacity, attention: level 3 (most of the time), break frequency: 4 hours, break length: 30 minutes, extreme break frequency: 4 hours and extreme break length: 30 minutes. This gives values for the shift timing component of 0.8 and 17.4 for the day and night shift respectively, and for the job type/breaks component values of 1.9 and 19.1, respectively. The cumulative component varies throughout the schedule. In Fig. 3 we show the output from the HSE tool for the Fatigue Index in (b) and the values for the three separate components in (c). On these panels we superimpose the values we obtain for the cumulative fatigue, 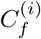 from equation (14) and for the Fatigue Index using our value for 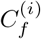 and the shift timing and job type components from the HSE tool.

## 2. The SAFTE model

### 2.1 Background

The Sleep, Activity, Fatigue, and Task Effectiveness (SAFTE) model describes the effectiveness of individuals based on their sleep history[5, 6]. The model was developed by members of the United States (US) of America’s Department of Defence (DOD) in the 1990s primarily as a way to model the effectiveness of members of the US armed forces during military missions but has since been used in other contexts [7].

Given a pattern of sleep and wake, the SAFTE model is designed to produce a measure of performance, termed ‘performance effectiveness’ which varies with time. There are several additional features available, including: ‘AUTOSLEEP’ tool that is designed to be used in conjunction with SAFTE to predict when sleep will occur on the basis of a shift pattern; a sleep fragmentation feature to model the effects of disrupted sleep; additional tools to calculate task specific scores. These are not discussed here.

The SAFTE model is described in [5] and closed source implementations exist such as the Fatigue Avoidance Scheduling Tool (FAST) [8]. The description below uses details from [5] and makes reasonable assumptions where details are missing, primarily around the rules for updating circadian phase.

Performance effectiveness *E*(%), is calculated as

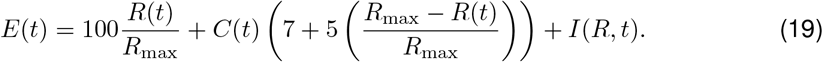

where *R*(*t*) is the level of a sleep reservoir that captures sleep history effects (sleep homeostasis), *C*(*t*) captures time of day (circadian) effects and *I*(*R, t*) is sleep inertia and models a short term reduction in effectiveness in the first 2 h after waking.

The performance effectiveness is capped above 100% and below 0%, conditions which do occur for some settings. A subject is at peak effectiveness when *E*(*t*) = 100%.

The following sub-sections describe the sleep reservoir, the circadian process and sleep inertia in more detail.

### 2.2 The sleep reservoir (a homeostatic process), *R*(*t*)

The homeostatic process is a measure of the physiological need for sleep. Within SAFTE, this is framed in terms of a sleep reservoir, the level of which is denoted by *R*(*t*) sleep units. While awake this reservoir is depleted at a rate of *K* = 0.5*/*60 sleep units s^*−*1^ down to a minimum of 0 units. Sleep replenishes the reservoir at a rate that is dependent on sleep intensity, where sleep intensity is greater when the reservoir is low and the reservoir is then replenished at a faster rate. The reservoir has a maximum capacity, *R*_max_ = 2880 sleep units.

Although not explicitly stated in this way in [5], obtaining outputs from the SAFTE model requires finding *R*(*t*) at time *t* by solving piecewise, non-autonomous ordinary differential equations:

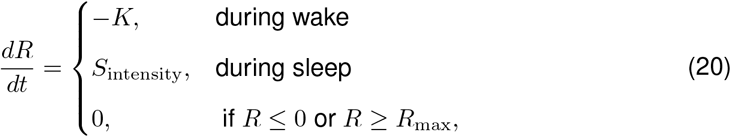

where *S*_intensity_ is sleep intensity and is modelled as

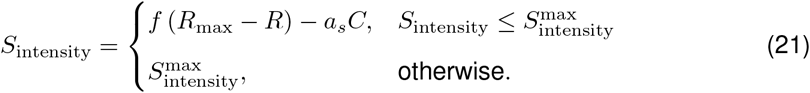

Here, *C*(*t*) is the time dependent circadian process discussed in the next subsection, *f* = 0.0026564*/*60 s^*−*1^, *a*_*s*_ = 0.55*/*60 s^*−*1^ and 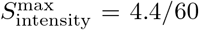 sleep units s^*−*1^. At time *t* = 0, the reservoir is assumed to be full so, *R*(0) = *R*_max_ sleep units. Note that in equations (20) and (21), for ease of reading we have dropped the explicit references to the variable dependencies.

An example of the behaviour of *R*(*t*) for one given sleep schedule is shown in Fig. 4(a). This shows the linear decrease in the level of the sleep reservoir with wake, with each hour of wake reducing the sleep reservoir by approximately 1%.

**Figure 4:**
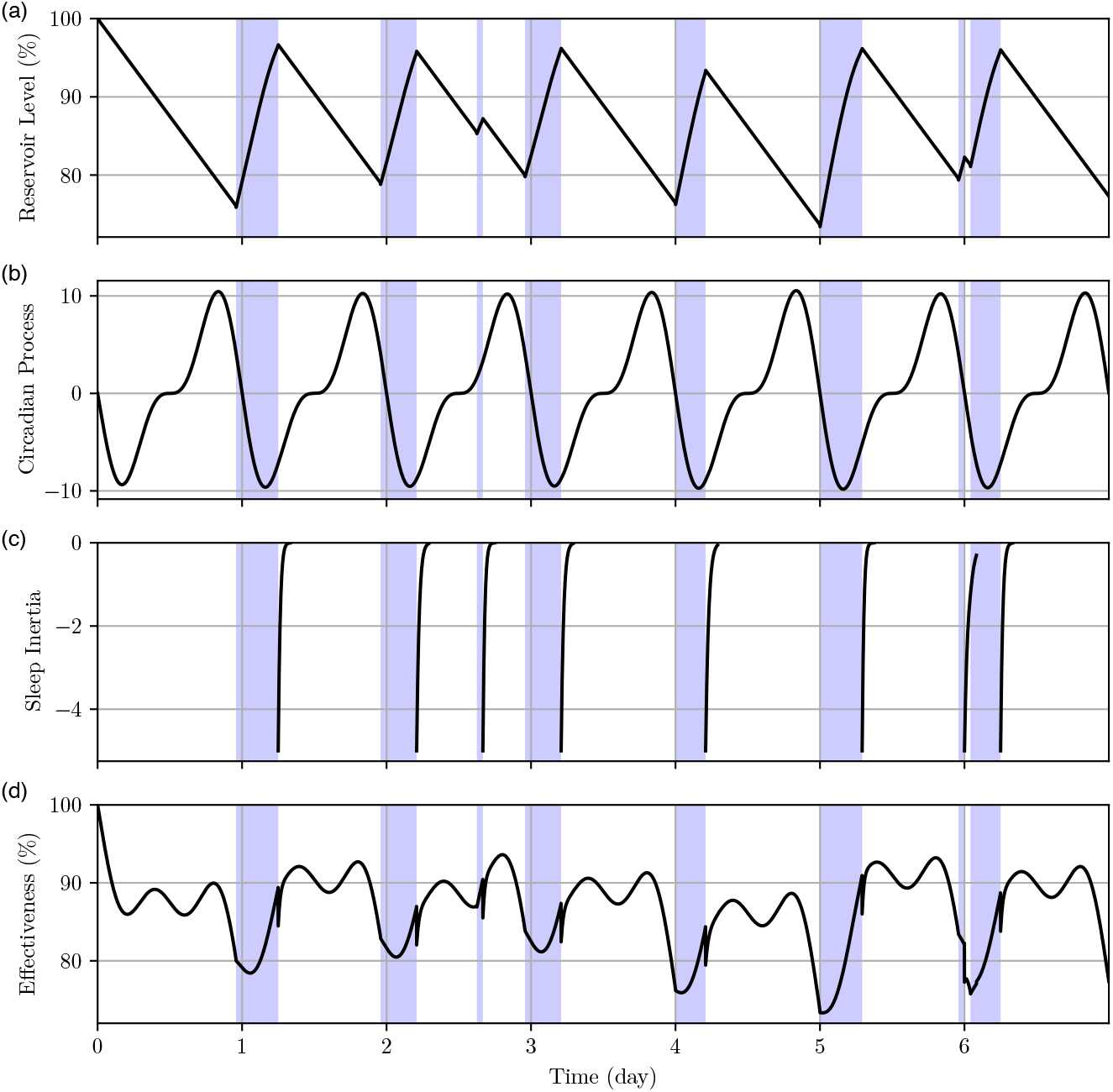
SAFTE: (a) The reservoir level *R*(*t*) as a function of time. The reservoir level drops during wake and rises during sleep. (b) The circadian process *C*(*t*) as a function of time. (c) The sleep inertia process *I*(*t*) as a function of time. Sleep inertia is non-zero only for 2 h immediately after waking. (d) The net performance effectiveness. For all panels, the intervals shaded as purple represent time intervals when sleep occurs.

### 2.3 Circadian process, *C*(*t*)

The circadian process is modelled as

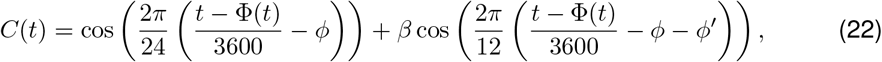

for *t* in seconds where the peak of the first curve is given as *φ* = 18 hours, the peak of the second curve is *ϕ* ^*′*^ = 3 hours from the first and the amplitude ratio between the two is given as *β* = 0.5 (unitless). The offset function Φ(*t*) is used to adjust for changes in circadian phase as a result of changes in a sleep pattern, such as during a time zone change or switching between day and night shifts. For situations with no change in circadian phase, Φ(*t*) is zero.

An example of the behaviour of *C*(*t*) for one given sleep schedule is shown in Fig. 4(b). When Φ(*t*) = 0, during the latter half of the normal working day *C*(*t*) is positive and counter-balances the depletion of the sleep reservoir. However, during normal sleeping hours *C*(*t*) is negative and reinforces the effects of the depleted sleep reservoir. In this way, *C*(*t*) helps maintain performance during the day but is an important factor in low levels of performance during the night.

There is little precise information on the equations for Φ(*t*) in [5]. Here we use a function that tries to match the description given as closely as possible. Since the equations are solved as discrete time steps, Φ(*t*) is generated as an iterative function at each of these time steps.

The offset at each time step in *t* is given as the previous offset plus an advance or delay, the choice of which depends on whether the individual’s current circadian phase matches their current sleeping pattern.

The phase shift for the first day is given as

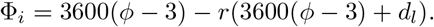

where *d*_*l*_ = 86400 is the day length in seconds and *r*(*t*) is the rolling average awake time. The rolling average is given by

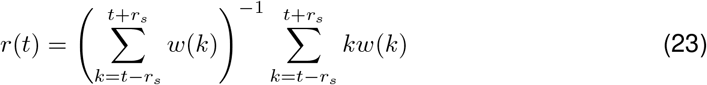

for *r*_*s*_ *≤ t < t*_*max*_ *− r*_*s*_, where *r*_*s*_ = 24 *×* 60 *×* 60*/*2 (s) is half the rolling window size, *t*_*max*_ = max(*t*) (s). When *t* = *r*_*s*_ or *t* = *t*_*max*_ *− r*_*s*_, *r*(*t*) is set to the first or last available value, respectively.

On subsequent days, the phase shift is

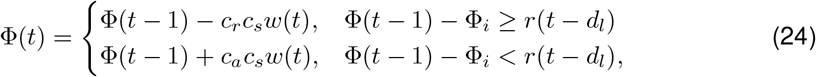

where *c*_*r*_ = 1*/*24 (unitless) is the circadian offset retreat rate, *c*_*a*_ = 1.5*/*24 (unitless) is the circadian offset advance rate and *c*_*s*_ = 24*/*18 (unitless) is a scaling factor. The function *w*(*t*) is 1 if the subject is awake at time *t* and 0 otherwise, and means that phase updates only occur while someone is awake.

### 2.4 Sleep inertia

Sleep inertia describes the observation that performance can be low for a short period after waking. In [5], this is captured by a sleep inertia variable *I* given by

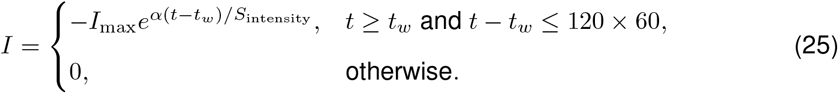

Here *I*_max_ = 5%, *α* = 0.04*/*3600 sleep units s^*−*2^ and *t*_*w*_ s is every time when the subject wakes up. The dependence of sleep inertia *I* on sleep intensity *S*_intensity_, means that sleep inertia is greater after waking from a higher intensity sleep. In addition, sleep inertia is assumed to only take effect for 0-2 h after waking. Note that in equation (25) for ease of reading we have dropped the explicit references to the variable dependencies.

An example of the behaviour of *I*(*t*) for one given sleep schedule is shown in Fig. 4(c), llustrating the performance loss of 5% immediately on waking and the dissipation of sleep intertia over the following 2 h.

### 2.5 Example: comparison with output from SAFTE

We did not have direct access to SAFTE. In order to test the equations in Section 2 we compared outputs from our model with outputs from SAFTE produced by Fatigue Science https://www.fatiguescience.com/.

As part of work commissioned by Dragados for TfL’s Crossrail project, Fatigue Science provided wristworn activity monitors (Readibands), guidance of their use for data collection and analysis of the subsequent data. The data collected included 30 days of actigraphy data for each of 30 tunnellers working a 7473 shift on Dragados’ TfL Crossrail project in 2014 and 30 days of actigraphy data for each of 35 tunnellers working a ‘day, back, night’ (DBN) shift on Dragados’ London Underground Bank Station Capacity Upgrade in 2018. Using sleep timings deduced from the actigraphy data, Fatigue Science used SAFTE to estimate fatigue levels. Fatigue Science then produced bar charts showing the average time spent at each performance effectiveness level. These bar charts are reproduced in Fig. 5(a) and Fig. 6(a) respectively.

**Figure 5:**
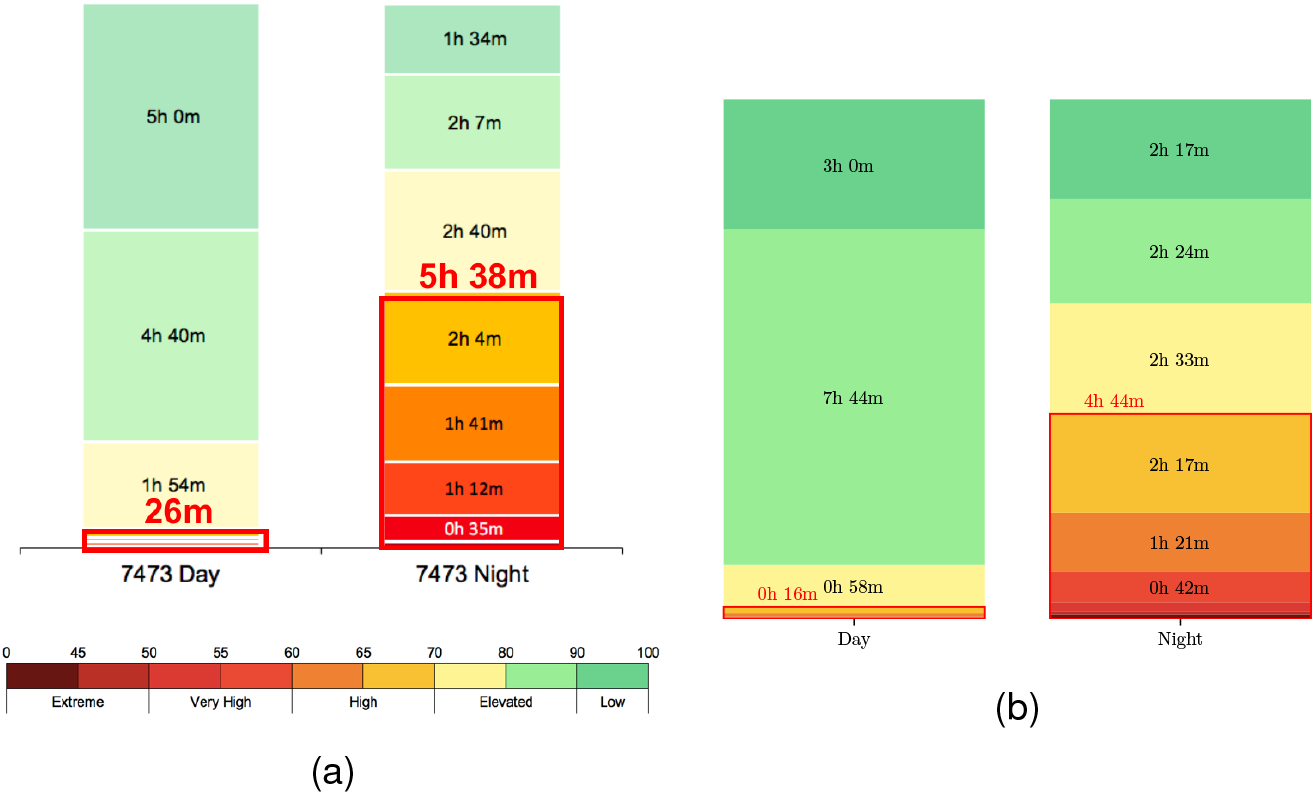
The average time spent at different performance effectiveness levels for the 7473 shift cycle. (a) Results from Fatigue Science. (b) Results as calculated from the equations in Section 2.

**Figure 6:**
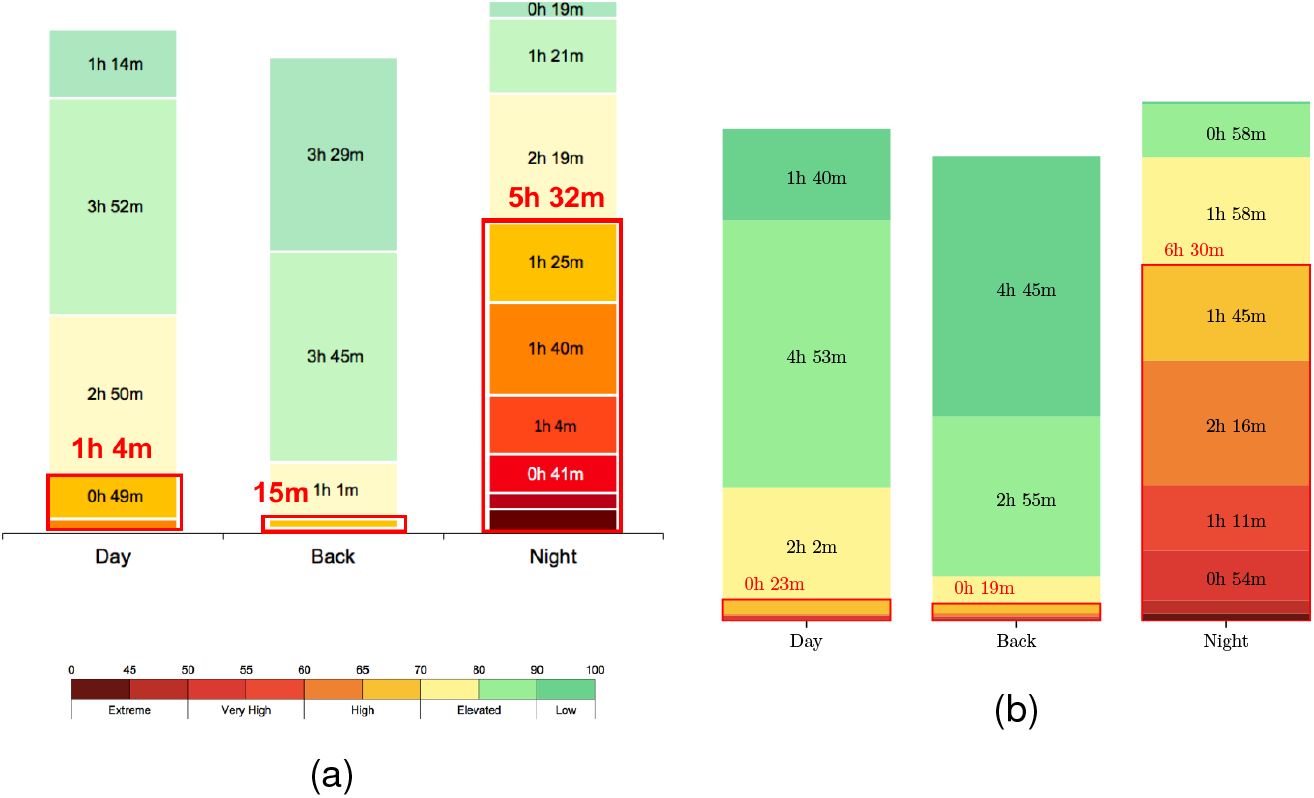
The average time spent at different performance effectiveness levels for the DBN shift cycle. (a) Results from Fatigue Science. (b) Results as calculated from the equations in Section 2.

We did not have full access to the actigraphy data, but we did have access to hourly aggregated data on activity for each worker and daily aggregated information on sleep duration. We used this aggregate information to estimate sleep timing then ran our SAFTE implementation for each worker and produced analogous charts to those made by Fatigue Science. Our bar charts for the 7473 and the DBN schedule are shown in Fig. 5(b) and Fig. 6(b) respectively.

Figures 5 and 6 show that both implementations are similar overall. For both implementations the day and back shifts had minimal time below 70% effectiveness, however night shifts have substantial time in this range with normally more than half the shift below 70%. In all instances the total time spend below 70% effectiveness was of the same magnitude and deviated at most by 58 minutes between implementations.

We believe that the differences are for two primary reasons. Firstly, we did not have access to the sleep timing inputs used by SAFTE. The aggregated data that were available limited our ability to reconstruct this sleep timing. We note that, within SAFTE, small differences in sleep timing can result in differences between adjacent categories (e.g. an increase in the 70-80 category and a decrease in the 80-90 category).

Secondly, the limited information available about the circadian shift calculation detailed in 2. This second reason is particularly relevant to explaining differences in the night shift, since it is the switch from the day shift to the night shift that results in a shift in the circadian rhythm. Consequently, differing implementations of the circadian shift calculation can drastically change the time spent below 70% effectiveness. We note that this is an area of considerable individual variability.

## 3 Comparison of the HSE Fatigue Index and SAFTE

Although we do not necessarily have the exact equations that either the HSE tool or the SAFTE model are based on, comparing the outputs of the equations we have stated here highlights the similarities and differences between the two approaches. We note that the outputs we show are outputs that it would be difficult to get from the tools in their current implementations.

The three cases discussed are: (i) the 7473 shift schedule for the ‘average’ sleeper; (ii) day-to-day variation in average sleep duration during a 7473 shift; (iii) the differences between a long and short sleeper undertaking a 7473 shift.

### 3.1 7473 shift schedules: ‘average’ sleepers

The 7473 shift schedule consists of seven day shifts of 12 h from 07:00 to 19:00, four days rest, seven night shift of 12 h from 19:00 to 07:00, followed by 3 days rest.

This shift schedule and results for the equations we have presented in this report for the HSE tool and for SAFTE are shown in Fig. 7 panels (a) and (b) respectively. Shift timing is indicated by the coloured horizontal bars, shaded according to the level of fatigue. SAFTE requires sleep timing as input, so in (b), the timing of sleep is shown by the grey bars. We have not implemented the extension of SAFTE, AUTOSLEEP, which estimates sleep timing on the basis of the shift schedule. Here, for the purpose of comparison with the HSE tool, we assume that sleep occurs from 23:00 to 05:15 on nights prior to day shifts and from 9:00 to 13:45 after night shifts. On rest days, we assume sleep occurs from 23:00 to 07:00. These timings were picked as representative of an ‘average’ sleeper who sleeps in accordance with approximate sleep durations from the HSE tool.

**Figure 7:**
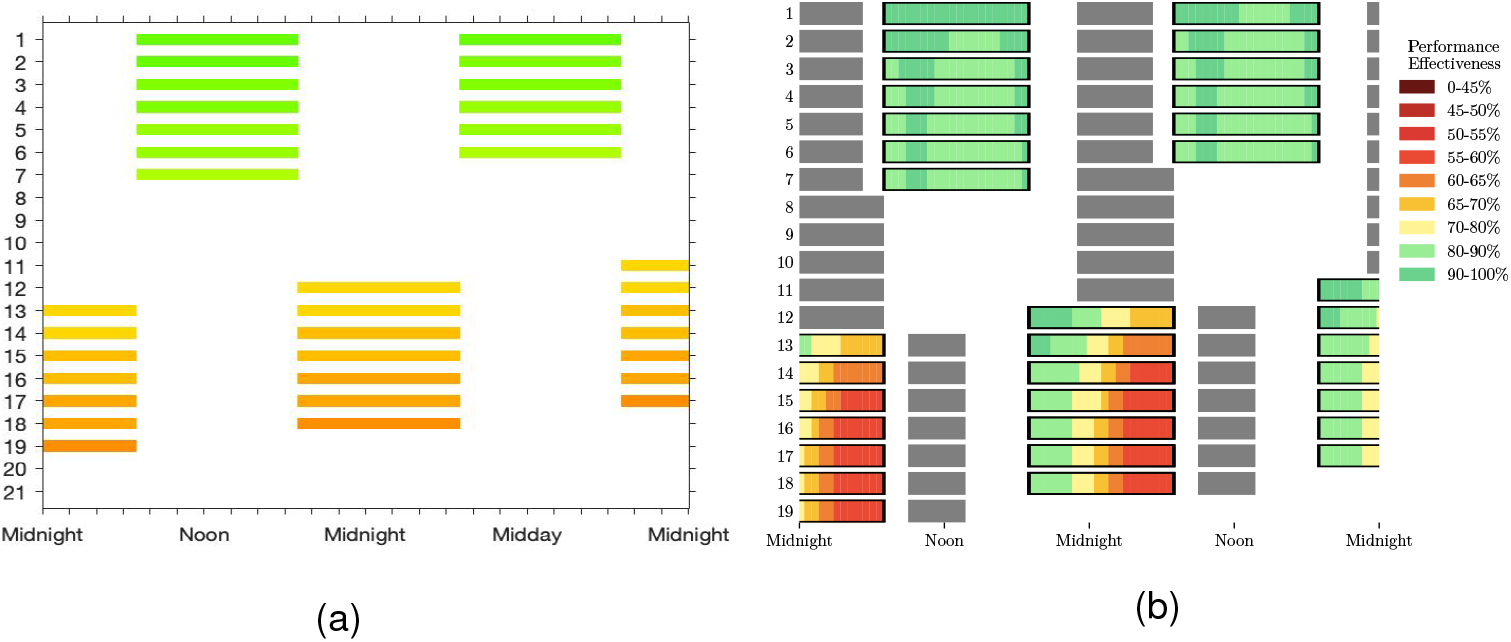
Comparison of fatigue predictions for the 7473 shift schedule. (a) The shift schedule with schedules shaded according to the *FI*^(*i*)^ (b) The equivalent output from SAFTE.

Successive days are plotted from top to bottom. The width of the plot covers two days, so that the top row shows the shift on day one followed by the shift on day two. On the next row, the shift on day two is re-plotted and then followed by the shift on day three. The reason for double-plotting in this way is so that both shifts that occur during the day and shifts that occur during the night can be seen clearly.

Fig. 7 illustrates two important points:

1. For an ‘average’ sleeper with similar assumptions on the amount of sleep obtained, both models give broadly the same message that during night shifts workers will feel more sleepy/perform less effectively.
2. SAFTE indicates the fatigue at different points of the shift. This is useful as it highlights that not all parts of the night shift are equal, with most problems likely to occur in the early hours of the morning.

The HSE tool does not output information on fatigue at different points during a shift, although in principle it is contained in the time of day component.

### 3.2 Day-to-day variation in sleep duration

Not surprisingly, the daily amount of sleep is fundamental to the determination of fatigue. Given a shift schedule, the HSE tool estimates how much sleep will occur in the breaks between successive shifts. Implicit in the data the HSE tool is based on is that this sleep occurs in one consolidated sleep episode.

The actigraphy data collected by Dragados and TfL in 2014, as described in Section 2.5, resulted in estimates for when and how much sleep actually occurred for the 30 tunnellers working a 7473 shift. Although actigraphy is known to over-estimate how much sleep occurs, it nevertheless provides a useful benchmark. In Table 1 we summarise the length of the sleep durations for each of the day, night and rest periods as estimated from our implementation of the HSE tool and from the activity monitors. From the activity data we have included results for the average across all tunnellers, and for the individiual with the longest sleep and the individual with the shortest sleep.

**Table 1:**
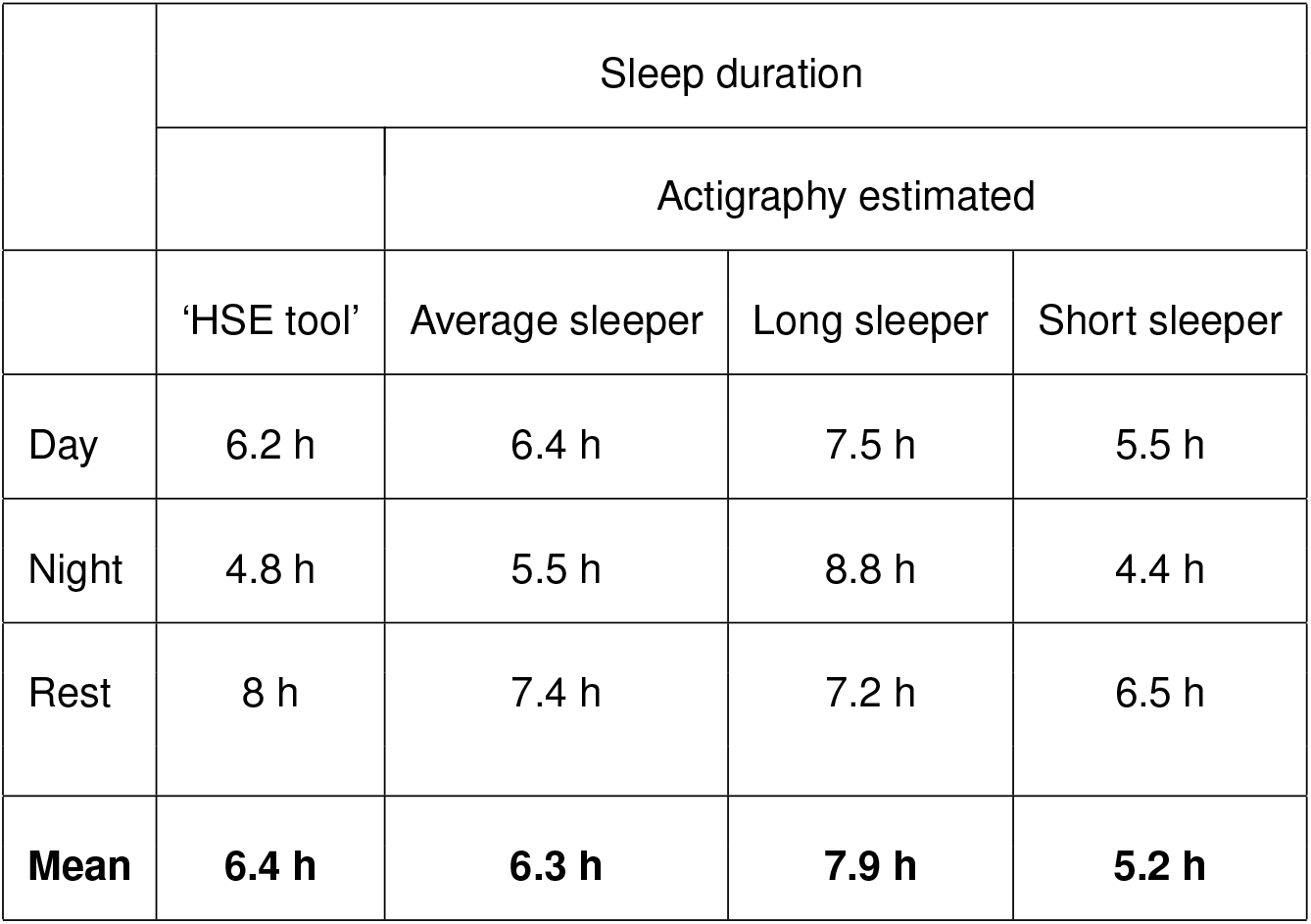
Comparison of sleep duration given by equation (4) for a 7473 shift schedule with sleep duration derived from actigraphy measures.

It can be seen that the HSE tool implementation compares well with the actigraphy estimates for the average amount of sleep. However, these averages hide some important day-to-day differences. In Fig.8 we compare the average for each day of the schedule for the HSE tool and from the actigraphy. Critically we see that there are substantial differences in the estimate of sleep duration at the transition from rest days to nights, with the HSE tool overestimating the amount of sleep.

We do not know what predictions the AUTOSLEEP function of SAFTE would make, but in the operational mode where SAFTE uses sleep timing derived from actigraphy SAFTE will be able to account for these pronounced day-to-day variations in a way that is not possible within the current HSE tool. In particular, since the HSE tool over-estimates sleep duration at the transition to night shifts, it could under-estimate fatigue for this period.

### 3.3 Long and short sleepers

No current commercial implementation of a biomathematical model of fatigue takes into account individual physiological differences in sleep need or preferred sleep timing. Personal circumstances are only taken into account through allowing different commute times.

The data from the Readiband study carried out by Fatigue Science for Dragados and TfL showed that, based on activity measures, there were pronounced differences between individuals in the amount of sleep they obtained (see Table 1). This is consistent with the known substantial differences between individuals [9].

Within current biomathematical models, differences in sleep duration then translate to different predictions for fatigue, as illustrated in Fig. 9 where outputs from our implementation of the HSE tool and for the SAFTE model are compared for the long and short sleeper. Since these models take no account of individual physiological differences, they necessarily predict that shorter sleep duration will result in more fatigue, as shown. Whether this is actually the case is not known: it could be that these two individuals differ in their sleep need.

**Figure 8:**
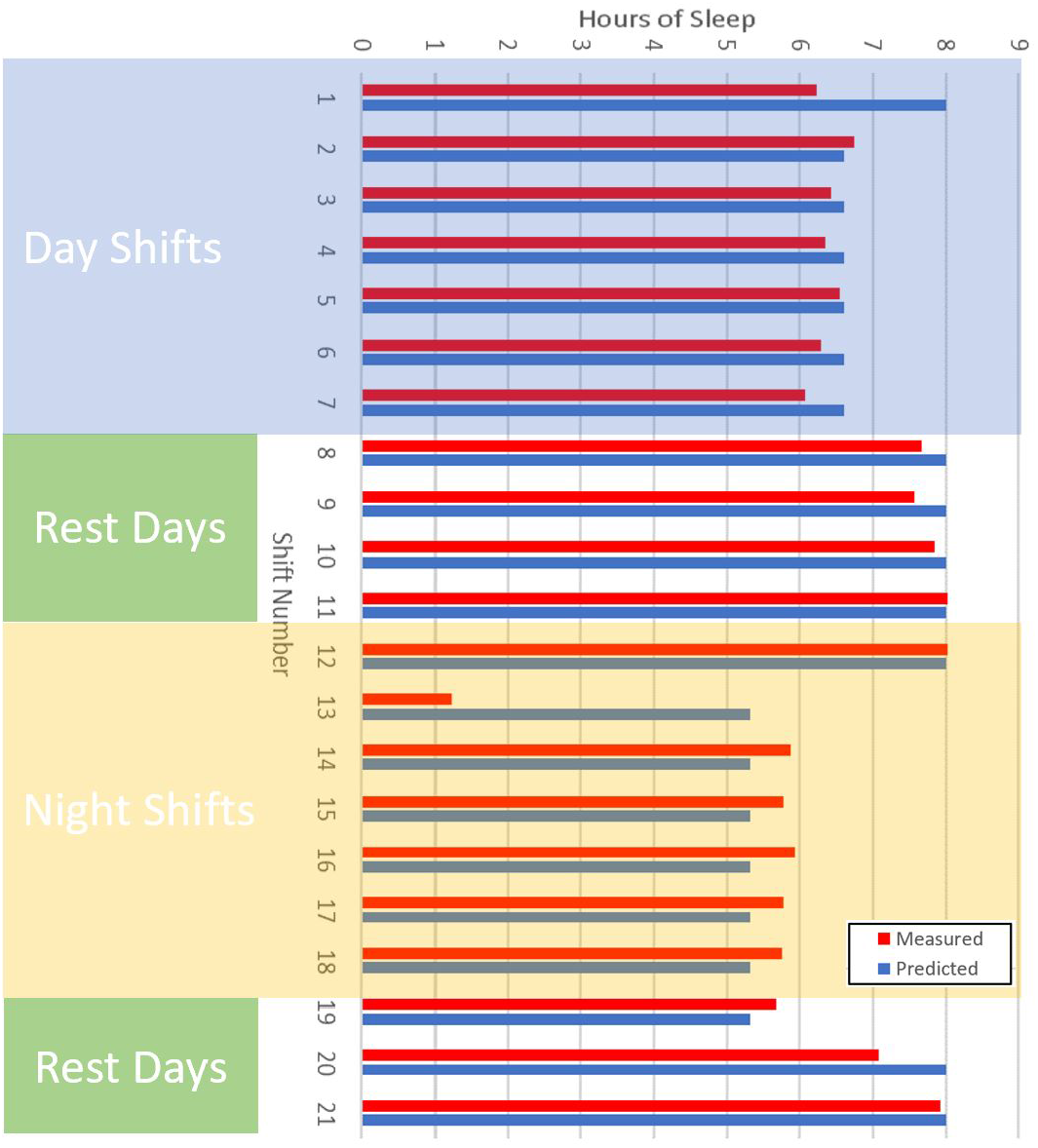
Comparison of average sleep durations across each day of the 7474 shift schedule as derived from actigraphy data versus predicted from equation (4) describing sleep duration as predicted by the HSE tool.

**Figure 9:**
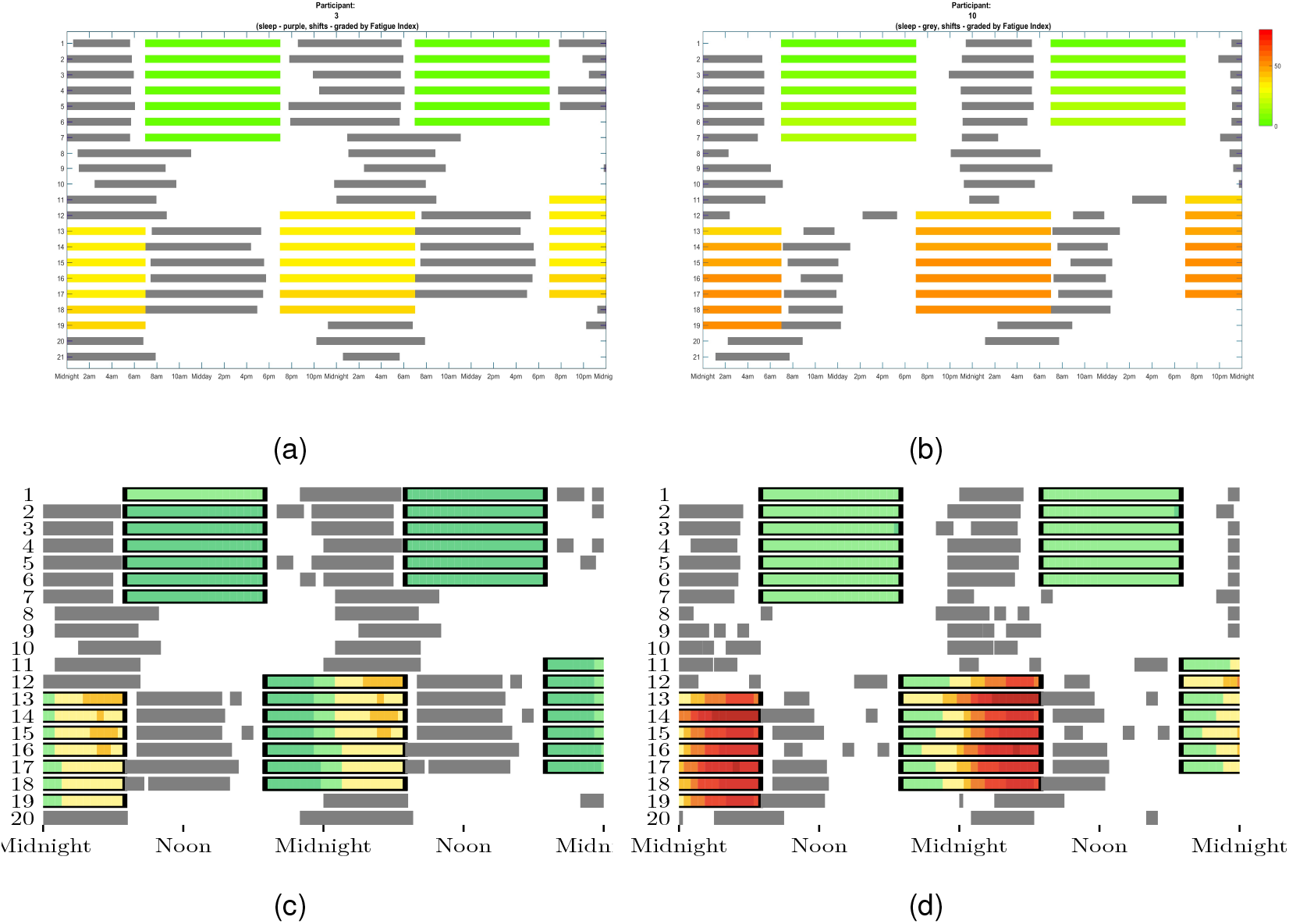
Short versus long sleepers: Predictions based on cumulated fatigue based on actual sleep using our model of the HSE tool (panels (a) and (b)) and our equations for SAFTE (panels (c) and (d)). Coloured bars indicate shift times and are coloured according to level of fatigue. Grey bars indicate the timing of sleep.

## 4 Concluding remarks

### 4.1 Data driven versus process driven models

All models are built on data.

The HSE tool is directly data-driven in the sense that the equations for the different componenents come from fitting to data that has been collected in a number of different experiments, some in the field and some in laboratory settings. For example: sleep loss due to early starts 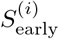 is based on the reported sleep times of Britannia crews on shifts ending at various times from the early evening through to mid-morning [10] and on the duration of sleep as a function of time of sleep onset for German and Japanese workers based on diary records, summarized by [11]. Sleep loss due to the pattern of shift was derived from the CHS studies of aircrew [12, 13, 10] and train drivers [14]; the shift timing components due to start time 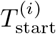, shift length 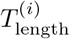 and time of day 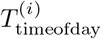 have been derived from several studies of aircrew and train drivers. The scalings from performance as measured by a psychomotor vigilance test to measures of sleepiness are based on laboratory studies [15, 16].

In contrast, SAFTE is process driven in the sense that data has been used to deduce that there are homeostatic and circadian processes and to fit parameters for these processes. There are pros and cons for each approach.

Both the HSE tool and SAFTE provide guidance on shift-scheduling with the user not required to input information on sleep duration. When operating in this mode, SAFTE needs input from AUTOSLEEP which estimates sleep durations. We did not have access to AUTOSLEEP so were unable to compare how sleep duration prediction compared with the HSE tool and therefore whether guidance from the two different models would differ. We did find that when similar assumptions were used about the daily sleep duration for a 7473 shift, the overall outputs of SAFTE and the HSE tool were similar.

An advantage of SAFTE is that it can produce estimates of fatigue risk broken down by hour whereas the HSE tool produces a single value for an entire shift. This is particular useful during night shifts, see Fig. 7 and Fig. 9, where an hour-by-hour break down highlights the fact that the beginning of a night shift and the end of a night shift are very different in terms of fatigue risk.

In addition, SAFTE has additional functionality to the HSE tool in that it can provide day-to-day guidance on who is currently fatigued and predict who will become fatigued over the next day. When operating in this mode, SAFTE takes sleep duration estimated from actigraphy bands.

### 4.2 Limitations of existing models

Some key missing elements in both of these models, and in fact all current commerically available biomathematical models of fatigue are listed below.

#### 4.2.1 Driving home

One significant risk of night shifts is the risk of accidents when driving home in the early hours of the morning. SAFTE can predict fatigue at all points of the day, although we understand that this is not typically provided as an output. For example, in Fig. 10 we have used our implementation of SAFTE to model fatigue for all waking hours for the long and short sleeper examples discussed in Section 3.3. The HSE model can, in principle, also predict fatigue at all points of the day, although it is not set up to do so at the moment.

**Figure 10:**
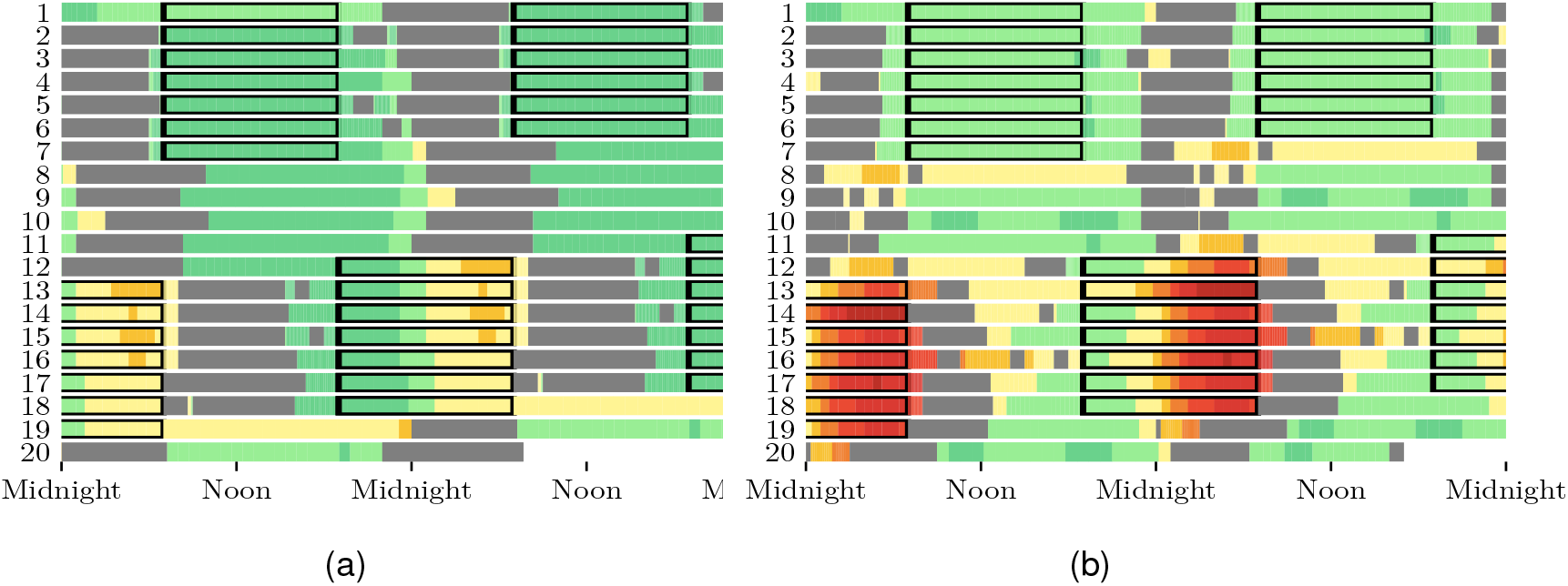
Fatigue for all points of the day. Here the grey bars indicate the sleep timing of the ‘short sleeper’ from the 7473 actigraphy study, the black boxes indicate the timing of the shift and the colours indicate levels of fatigue. This is the same data as shown in Fig. 9(c) and (d) but with fatigue also calculated for times that the individual is not on shift.

#### 4.2.2 Starting well-rested

All models have to make some assumption about the level of fatigue at the start of the schedule. Both the HSE tool and SAFTE make an assumption that someone starts their shifts well-rested. Personal circumstances and choices can mean that this is not the case. In fact, an important element of shift design is careful choice of the start time for the first shift in a schedule.

#### 4.2.3 Individual differences

There are large individual differences in sleep preference and sleep need [9]. Neither the HSE tool nor the SAFTE model are currently able to model individual physiological differences. As seen in Fig. 9 they predict that someone with a shorter sleep duration will necessarily be more fatigued than someone with a longer sleep duration. This may not be the case. While there is research that has considered tuning models for individuals, for example, [17, 18, 19], an operational model that includes uncertainty due to individual differences is not yet available.

#### 4.2.4 The light environment

Light has a short term alerting effect such that the colour and intensity of light during night shifts can affect performance. Light also has longer term effects on the biological clock and preferred sleep timing. There has been much recent work on developing recommendations for healthy light environments [20]. Mathematical models that include the impact of light on biological timing have been constructed [21, 22, 23] but are not yet part of operational models.

#### 4.2.5 Transparency

Commercial sensitivities mean that most models are not transparent with many of the assumptions they are built on not publicly available. This makes them difficult to independently verify. Mathematical models of sleep can have complex behaviour [24, 25] and it is not clear that this is fully recognised in their implementation. It is also difficult to make direct comparisons between models. For example, with the HSE tool and SAFTE they take different inputs and output different scales.

#### 4.2.6 Translation of outputs to meaningful workplace outcomes

The outputs of some models are hard to relate to quantities that are measureable in the workplace.

### 4.3 Future directions

#### 4.3.1 Guidance on scheduling and education on sleep and fatigue should be considered at least as important as current biomathematical models

The core physiological principles that describe fatigue due to lack of sleep are that fatigue is dependent on (i) sleep history: the longer that you have been awake, the greater your physiological need for sleep; sleep debt accumulates across days and weeks; (ii) time of day effects: across a 24 h period, fatigue oscillates according to an internal physiological clock (the circadian clock).

In their current form, it is not clear that biomathematical models can provide more information than an expert with a firm grasp of these twin concepts of sleep history and time-of-day effects. The role of biomathematical models is then a way of providing the non-expert with a tool to manage fatigue. But use of models without understanding their scope and limitations is itself risky. Therefore, guidance on scheduling and education on sleep and fatigue should be considered at least as important as current biomathematical models.

#### 4.3.2 Only by analysing and integrating high quality individual data on sleep, fatigue, performance, near misses, accidents, actual shift patterns with models can we develop better models and management systems to reduce fatigue and associated risks

With new technological solutions including wearables, nearables and apps there is huge scope to collect more high quality data in order to improve biomathematical models of fatigue. Note while it is useful to collect measures of sleep via actigraphy, collecting outcome measures such as sleepiness and performance is critical. For example, current models predict that individuals who sleep less will be at a greater fatigue risk than those who sleep more. However, those in their 50’s typically sleep less than those in their 20’s [26], but there is little evidence that they are sleepier. In fact the evidence is rather the reverse [27]. As further illustration, recent studies have highlighted that different people working on the same shift pattern have different sleep patterns [28] but without knowing the consequent impact on individual cognitive function, it is hard to use this information to refine biomathematical models. Models can account for individual differences in, for example, sleep need [29] and the effect of light [23], but are not yet part of commercial biomathematical models of fatigue.

#### 4.3.3 Individual self-management of fatigue

The importance of making time for sleep is not always recognised. Education, early diagnosis of sleep disorders such as sleep apnea, and self-monitoring all have a role to play in reducing fatigue-related risk in the work-place.

Self-monitoring could be via the many actigraphy-based sleep trackers available. But as with the use biomathematical models, care is needed [30]. Wearables cannot reliably capture differences between sleep and sitting quietly. Once in bed, actigraphy is good at identifying sleep, but periods of quiet wakefulness are frequently mis-classified as sleep. For example, the Readiband used by Fatigue Science is reported as having a sensitivity of 88% and a specificity of 55% for detecting sleep while in bed [31]. This means that 88% of the time sleep occured the actigraphy algorithm correctly identified sleep. However, 45% of the time that wake occurred is was also classified as sleep.

Mis-information on sleep duration can result in actual differences in performance [32] and promote anxiety over sleep.

The passive nature of sleep trackers, with no need for input from the wearer is a positive. A more active approach where individuals enter and track their own subjective measures on sleep quality and measures of wellbeing could be an alternative [33].

#### 4.3.4 Final comments

From publicly available data, it appears the mathematical models that underpin the HSE tool and those that underpin SAFTE have not been updated in well over a decade. Some of the other commercially available models also appear to have undergone little change in the underlying mathematics, with most development focussing on the (necessary) design of interfaces and applying models in the field.

While the integration with wearables is an important step, and appropriate interfaces absolutely essential, there is considerable scope for the further development of the underlying mathematical models.

Combining high quality data with improvements to mathematical models could result in better fatigue risk management tools that capture not only risk but also uncertainty in risk predictions due to individual factors.

## 5 Definition of symbols

### HSE model

HSE model:
|  |  |
| --- | --- |
| $FI^{(i)}$ | Fatigue Index for day $i$ |
| $C_f^{(i)}$ | Cumulative fatigue component of Fatigue Index on day $i$ |
| $\bar{T}_f^{(i)}$ | Shift timing component of Fatigue Index on day $i$ |
| $\bar{J}_f^{(i)}$ | Job type component of Fatigue Index on day $i$ |
| $S_{\text{early}}^{(i)}$ | Predicted sleep duration in hours on day $i$ as a result of an early start |
| $S_{\text{bed}}^{(i)}$ | Predicted sleep duration in hours on day $i$ for a given bedtime |
| $S_{\text{late}}^{(i)}$ | Predicted sleep duration in hours on day $i$ as a result of a late finish |
| $S_{\text{quick}}^{(i)}$ | Predicted sleep duration in hours on day $i$ as a result of a quick return |
| $S_{\text{duration}}^{(i)}$ | Resultant sleep duration in hours on day $i$ |
| $S_{\text{loss}}^{(i)}$ | Sleep loss in hours on day $i$ |
| $P_{\text{baseline}}^{(i)}$ | Baseline performance as measured by the PVT on day $i$ for the given sleep loss |
| $P_{\text{cumulative}}^{(i)}$ | Performance $i$ as a result of accumulated sleep loss |
| $S_p^{(i)}$ | Predicted sleepiness on the Samn Perelli scale |
| $S_{pcc}^{(i)}$ | |
| $pKSS^{(i)}$ | |
| $t$ | Time |
| $t_{\text{start}}^{(i)}$ | Shift start time shift $i$ |
| $t_{\text{bed}}^{(i)}$ | Bedtime for shift $i$ |
| $t_{\text{finish}}^{(i)}$ | Shift finish time for shift $i$ |
| $t_{\text{gap}}^{(i)}$ | Gap between shift on day $i$ and the shift on day $i - 1$ |
| $t_{\text{commute}}^{(i)}$ | Total time spent commuting on day $i$. |
| $T_{\text{start}}^{(i)}$ | Contribution to shift timing component of start time |
| $T_{\text{length}}^{(i)}$ | Contribution to shift timing component of the length of the shift |
| $T_{\text{timeofday}}^{(i)}$ | Contribution to shift timing component of the time of day of the shift |

### SAFTE model

**SAFTE model:**
|  |  |
| --- | --- |
| $E(t)$ (%) | Performance effectiveness |
| $R(t)$ | Reservoir level |
| $C(t)$ | Circadian rhythm |
| $I(R, t)$ | Sleep inertia |
| $t$ | Time |
| $R_{\max} = 2880$ units | Reservoir capacity |
| $S_{\text{intensity}}$ sleep units $\text{s}^{-1}$ | Sleep intensity |
| $S_{\text{intensity}}^{\max} = 4.4/60$ sleep units $\text{s}^{-1}$ | Maximum sleep intensity |
| $K = 0.5/60\text{s}^{-1}$ | Rate of reservoir depletion |
| $f = 0.0026564/50\text{s}^{-1}$ | |
| $a_s = 0.55/60\text{s}^{-1}$ | a weighting factor |
| $\phi = 18 : 00$ | Time of the maximum of the 24 h rhythm |
| $\phi' = 3 : 00$ | Time of the maximum of the 12 h rhythm relative to the 24 h rhythm |
| $\Phi$ | Shift of the circadian rhythm due to changes in sleep timing |
| $\beta = 0.5$ | Amplitude of the 12 h rhythm relative to the 24 h rhythm |
| $I_{\max} = 5\%$ | maximal inertial effect on awakening |
| $\alpha = 0.04/60$ sleep units $\text{s}^{-2}$ | inertia time constant |

## Footnotes

1 Crossrail is the biggest railway infrastructure project in Europe and is one of the largest single investments undertaken in the UK. It is a joint venture between TfL and the UK Department of Transport. Construction started in 2009 and intensive operational testing is expected to take place in 2021. The project included digging 42 kilometres of new rail tunnel under London. Dragados was one of two engineering companies responsible for tunnelling.

2 or ‘duty’

3 A technical point: we have used the index *i* to denote the day on which the shift occurs. More formally, we should include both an index for the day and for the shift number to allow for the fact that more than one shift can occur on one day and there may not be a shift on every day. However, to keep the notation from being overly cumbersome, for the description here we have only specified a single index for the day. In any computer coding of the model, appropriate care has to be taken for both days and shifts.

4 The values reported in the Fatigue Index spreadsheet in the columns labelled ‘cumulative fatigue’, ‘duty timing’ and ‘duty type’ are 100 times the values for 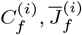 and 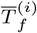

5 For coding purposes, for any shift it is important to determine which of [ineq005D and 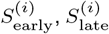 and 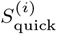 apply.

